# Angiotensin II Inhibits the A-type K^+^ Current of Hypothalamic Paraventricular Nucleus Neurons in Rats with Heart Failure: Role of MAPK-ERK1/2 Signaling

**DOI:** 10.1101/2021.09.12.459939

**Authors:** Ranjan K. Roy, Hildebrando Candido Ferreira-Neto, Robert B. Felder, Javier E. Stern

## Abstract

ANGII-mediated sympathohumoral activation constitutes a key pathophysiological mechanism in heart failure (HF). While the hypothalamic paraventricular nucleus (PVN) is recognized as a major site mediating ANGII effects in HF, the precise mechanisms by which ANGII influences sympathohumoral outflow from the PVN remain unknown. ANGII activates the ubiquitous intracellular MAPK signaling cascades and recent studies revealed a key role for ERK1/2 MAPK signaling in ANGII-mediated sympathoexcitation in HF rats. Importantly, ERK1/2 was reported to inhibit the transient outward potassium current (I_A_) in hippocampal neurons. Given that I_A_ is a critical determinant of the PVN neuronal excitability, and that downregulation of I_A_ in the brain has been reported in cardiovascular disease states, including HF, we investigated here whether ANGII modulates I_A_ in PVN neurons via the MAPK-ERK pathway, and, whether these effects are altered in HF rats. Patch-clamp recordings from identified magnocellular neurosecretory (MNNs) and presympathetic (PS) PVN neurons revealed that ANGII inhibited I_A_ in both PVN neuronal types, both in sham and HF rats. Importantly, ANGII effects were blocked by inhibiting MAPK-ERK signaling as well as by inhibiting EGFR, a gateway to MAPK-ERK signaling. While no differences in basal I_A_ magnitude were found between sham and HF rats under normal conditions, MAPK-ERK blockade resulted in significantly larger I_A_ in both PVN neuronal types in HF rats. Taken together, our studies show that ANGII-induced ERK1/2 activity inhibits I_A_ and increases the excitability of presympathetic and neuroendocrine PVN neurons, contributing to the neurohumoral overactivity than promotes progression of the HF syndrome.

**GRAPHICAL ABSTRACT:** 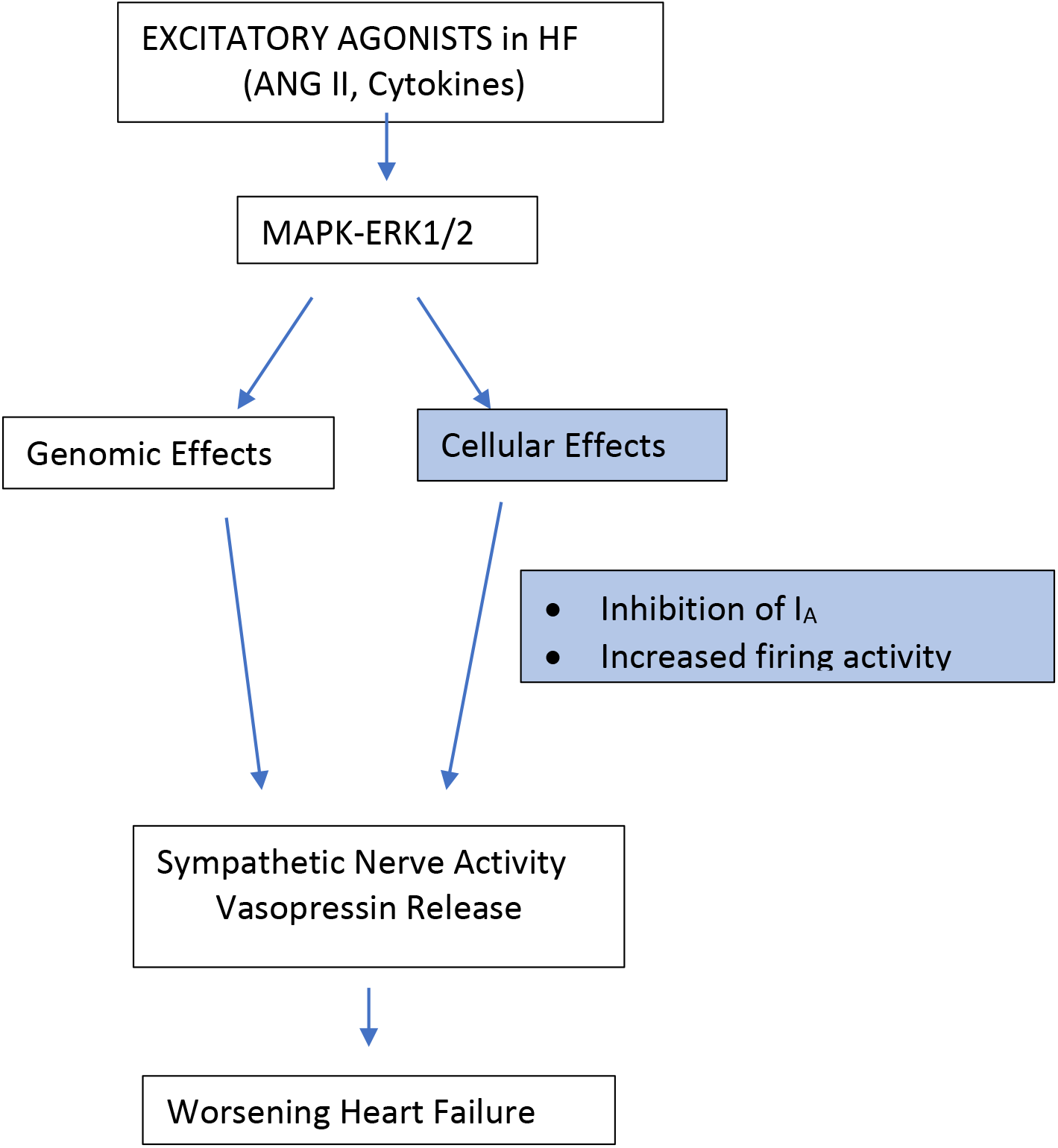

## INTRODUCTION

The renin-angiotensin system (RAS), in particular the peptide angiotensin II (ANGII), plays critical roles in the maintenance of fluid and cardiovascular homeostasis. In addition to its peripheral actions, ANGII acts within the central nervous system (CNS) to stimulate autonomic and neuroendocrine outputs to peripheral tissues (1). Importantly, ANGII-mediated sympathohumoral activation constitutes a key pathophysiological mechanism in neurogenic hypertension and heart failure (HF) (2–7). Despite the importance of central ANGII actions, the precise mechanisms underlying ANGII-mediated sympathetic activation remain unknown.

Within the CNS, the paraventricular nucleus of the hypothalamus (PVN) is a major substrate mediating central ANGII action (8–10), and an increase in PVN RAS activity is known to contribute to sympathoexcitation in HF (5, 11–13). Thus, interventions that reduce RAS activity in the PVN diminish sympathetic outflow and improve the systemic manifestations of HF (11, 14–17).

Neurohumoral outflow from the PVN is largely determined by the activity of both magnocellular neurosecretory and presympathetic neurons that project to the rostral ventrolateral medulla (RVLM) in the brainstem and to preganglionic sympathetic neurons in the spinal cord. Importantly, both *in vivo* (8, 18, 19) and *in vitro* (9, 20–23) electrophysiological studies have demonstrated that ANGII increases the discharge frequency of both PVN neuronal types. Still, the precise signaling mechanisms by which ANGII influences PVN neuronal activity and sympathohumoral outflow, both in health and disease conditions, remains to be determined.

ANGII is known to activate the ubiquitous intracellular MAPK signaling cascades (24–27), and recent studies have revealed a key role for brain ERK1/2 MAPK signaling in ANGII-mediated sympathetic excitation in HF rats (13, 28). Moreover, we recently showed that inhibition of brain MAPK signaling, significantly reduces sympathetic nerve activity in HF rats (13, 28). Taken together, these studies support the notion that ANGII signaling via the MAPK-ERK pathway constitutes a key signaling mechanism contributing to neurohumoral activation in HF.

Phosphorylation of ERK1/2 ultimately elicits transcription of proteins that may sustain the excitatory neurochemical milieu of the PVN. Thus, studies examining the effects of chronic interference with ERK1/2 signaling in HF rats have revealed reductions in brain renin-angiotensin system components (13, 28) inflammatory mediators (13, 28) and indicators of endoplasmic reticulum stress (13). Since all of these have been shown to contribute to the excessive sympathetic drive in HF, it may be presumed that these genomic effects of ERK1/2 contribute to the sympathoexcitatory influence of ERK1/2 signaling. However, the acute effects of ERK1/2 inhibition, which has been shown to rapidly reduce blood pressure and sympathetic outflow in rats with established HF (13) and to minimize pressor responses to excitatory agonists in normal rats (13, 29), occur far too early to be explained by protein synthesis and suggest a more direct effect on the excitability of PVN neurons. Of particular interest in this regard, ERK1/2 has been reported to phosphorylate and inactivate the voltage gated potassium channel (Kv) subunit Kv4.2 in hippocampal neurons (30–32), reducing the transient outward potassium current (I_A_). Phosphorylation of Kv4.2 occurs rather rapidly, not requiring gene transcription. Kv4.2 is also expressed in PVN neurons (33, 34), and I_A_ is an important determinant of the excitability of both magnocellular and parvocellular PVN neurons (34–37). Moreover, we have previously shown that down regulation of I_A_ availability contributes to increased hypothalamic neuronal excitability during hypertension (37).

The present study investigated whether ANGII modulates I_A_ in magnocellular and presympathetic PVN neurons, whether these actions involve the MAPK-ERK pathway, and finally, whether these effects are altered in rats with HF. We hypothesized than ANGII-induced ERK1/2 activity increases the excitability of presympathetic and neuroendocrine PVN neurons, contributing to the neurohumoral overactivity than promotes progression of the HF syndrome.

## MATERIALS AND METHODS

### Animals

Male Wistar rats purchased from Envigo (Harlan) (4-6 weeks old) were housed under standardized conditions (12:12 h light–dark cycle, lights on 07.00 h) with food and water available *ad libitum*. All experimental procedures were in strict compliance with NIH guidelines, and were approved by the Georgia State University Institutional Animal Care and Use Committee.

### Induction of heart failure (HF)

HF was induced by coronary artery ligation as described previously (38, 39). Briefly, animals were anesthetized with isoflurane (2-4%) and intubated for mechanical ventilation. A left thoracotomy was performed, and the heart exteriorized. The ligation was placed on the left anterior descending coronary artery. Buprenorphine SR-LAB (0.5 mg/kg sc; Zoo Pharm, Windsor, CO, USA) was given immediately after surgery to minimize postsurgical pain. Sham control animals underwent the same procedure with the exception that the coronary artery was not occluded. Transthoracic echocardiography (Vevo 3100 systems, Visual Sonics, Toronto, ON, Canada) was performed 4 weeks after surgery under light isoflurane (2-4%) anesthesia. The measurements of the left ventricle internal diameter and its posterior and anterior walls in systole and diastole, and the ejection fraction and fractional shortening were obtained throughout the cardiac cycle from the short-axis motion imaging mode. The mean ejection fraction (EF) in sham (n=16) and HF (n=14) rats were 89.3 ± 0.6 and 33.1 ± 1.1, respectively (p< 0.0001, unpaired t test).

### Identification of presympathetic PVN neurons

Presympathetic RVLM-projecting PVN neurons were identified by a combination of retrograde tracing and electrophysiological characteristics. For retrograde tracing, rhodamine beads were injected unilaterally into the brainstem region containing the RVLM as previously described (34). Rats were anesthetized with isoflurane (2-4%) and a stereotaxic apparatus was used to pressure inject 200 nl of rhodamine-labeled microspheres (Lumaflor, Naples, FL, USA) into the RVLM (starting from Bregma: 12 mm caudal along the lamina, 2 mm medial lateral, and 8 mm ventral). As previously reported, RVLM injection sites were contained within an area from the caudal pole of the facial nucleus to ~ 1 mm more caudal, and were ventrally located with respect to the nucleus ambiguus. The location of the tracer was subsequently verified histologically (34). Rats were used 2-3 days after surgery. The recorded neurons displayed membrane properties characteristic of parvocellular presympathetic neurons, namely the presence of a low-threshold spike and absence of a transient outward rectification (36, 40, 41).

### Hypothalamic slices

Rats were anaesthetized with pentobarbital (50 mg kg-1), quickly decapitated, and brains were dissected out. Coronal slices were cut (250 μm thick) utilizing a vibroslicer (Leica VT1000). An oxygenated ice cold artificial cerebrospinal fluid (ACSF) was used during slicing (containing in mM): 119 NaCl, 2.5 KCl, 1 MgSO_4_, 26 NaHCO_3_, 1.25 NaH_2_PO_4_, 20 D-glucose, 0.4 ascorbic acid, 2.0 CaCl_2_ and 2.0 pyruvic acid; pH 7.4; 290–310 mOsmol/l?). After sectioning, slices were placed in a holding chamber containing ACSF and kept at room temperature (22°C) until used.

### Drugs

Tetrodotoxin (TTX) and kynurenic acid were purchased from Alomone Laboratories (Jerusalem, Israel). Tyrphostin and PD98059 were purchased from ThermoFisher Scientific (Pittsburgh, Pennsylvania).

### Electrophysiological recordings

Slices were placed in a submersion style recording chamber, bathed with solutions (3.0ml/min^-1^) that were bubbled continuously with a gas mix of 95% O_2_–5% CO_2_, and maintained at near physiological temperature (32°C). Thin-walled (1.5 mm o.d., 1.17 mm i.d.) borosilicate glass (G150TF-3, Warner Instruments, Sarasota, FL, USA) was used to pull patch pipettes (3–6MΩ) on a horizontal Flaming/Brown micropipette puller (P-97, Sutter Instruments, Novato, CA, USA). PVN neurons were selected for whole-cell patch-clamp recordings using DIC videomicroscopy and epifluorescence (retrogradely-labeled neurons). Recordings were obtained with a Multiclamp 700A amplifier (Axon Instruments, Union City, CA, USA). The voltage output was digitized at 16-bit resolution, 10 kHz (Digidata1320A, Axon Instruments), and saved on a computer to be analyzed offline using pCLAMP9 software (Axon Instruments). The experiment was discarded if the series resistance was unstable (changes >20%). The internal solution contained (in mM): 140 potassium gluconate, 0.2 EGTA, 10 HEPES, 10 KCl, 0.9 MgCl_2_, 4 MgATP, 0.3 NaGTP and 20 phosphocreatine (Na^+^); pH 7.2–7.3. _I_A_ current density was determined by dividing the current amplitude by the cell capacitance, obtained by integrating the area under the transient capacitive phase of a 5 mV depolarizing step pulse. I_A_ was activated using conventional protocols (depolarizing steps from −70 to +10 mV in 10 mV, 300 ms) (37). I_A_ was isolated electronically by digital subtraction of currents evoked from two separate protocols, in which preconditioning steps of either −120 mV and −10 mV were used, to fully evoke, and to completely inactivate I_A_, respectively (42). Since the focus of present study was on I_A_, the properties of I_KDR_ were not further studied.

### Statistical Analysis

All values are expressed as means ± SEM. Student’s paired *t* test was used to compare the effects of a drug treatment on I_A_. Between group differences values were compared using analysis of variance repeated measures (ANOVA-RM). Where the *F* ratio was significant, *post hoc* comparisons were completed using the Bonferroni *post hoc* test. Differences were considered statistically significant at *P*<0.05 and *n* refers to the number of cells. All statistical analyses were conducted using GraphPad Prism (GraphPad Software, San Diego, CA, USA).

## RESULTS

### ANGII inhibits the magnitude of I_A_ in magnocellular neurosecretory and presympathetic PVN neurons

Whole-cell patch clamp recordings were obtained from hypothalamic PVN neurons that based on their size, electrophysiological properties and retrograde labeling (see Methods) were identified either as magnocellular neurosecretory neurons (MNNs, n=71) or presympathetic neurons (PSs, n=66). Bath application of ANGII (0.5 μM, 8 minutes ***Fig.1***) evoked a progressive inhibition of I_A_ magnitude both in MNNs (F= 8.04, p< 0.0001, One way ANOVA, n= 15) as well as in PS (F= 4.02, p< 0.0001, One way ANOVA, n= 19). Interestingly, whereas no recovery in I_A_ magnitude was observed after washout in MNNs, a partial recovery was observed in PSs. No differences in the peak of ANGII-mediated inhibition of I_A_ was observed between the two cell types (−19.7 ± 3.2 % vs −21.12 ± 5.0 % in MNNs and PSs respectively, P= 0.81, unpaired t test).

**Fig 1.**
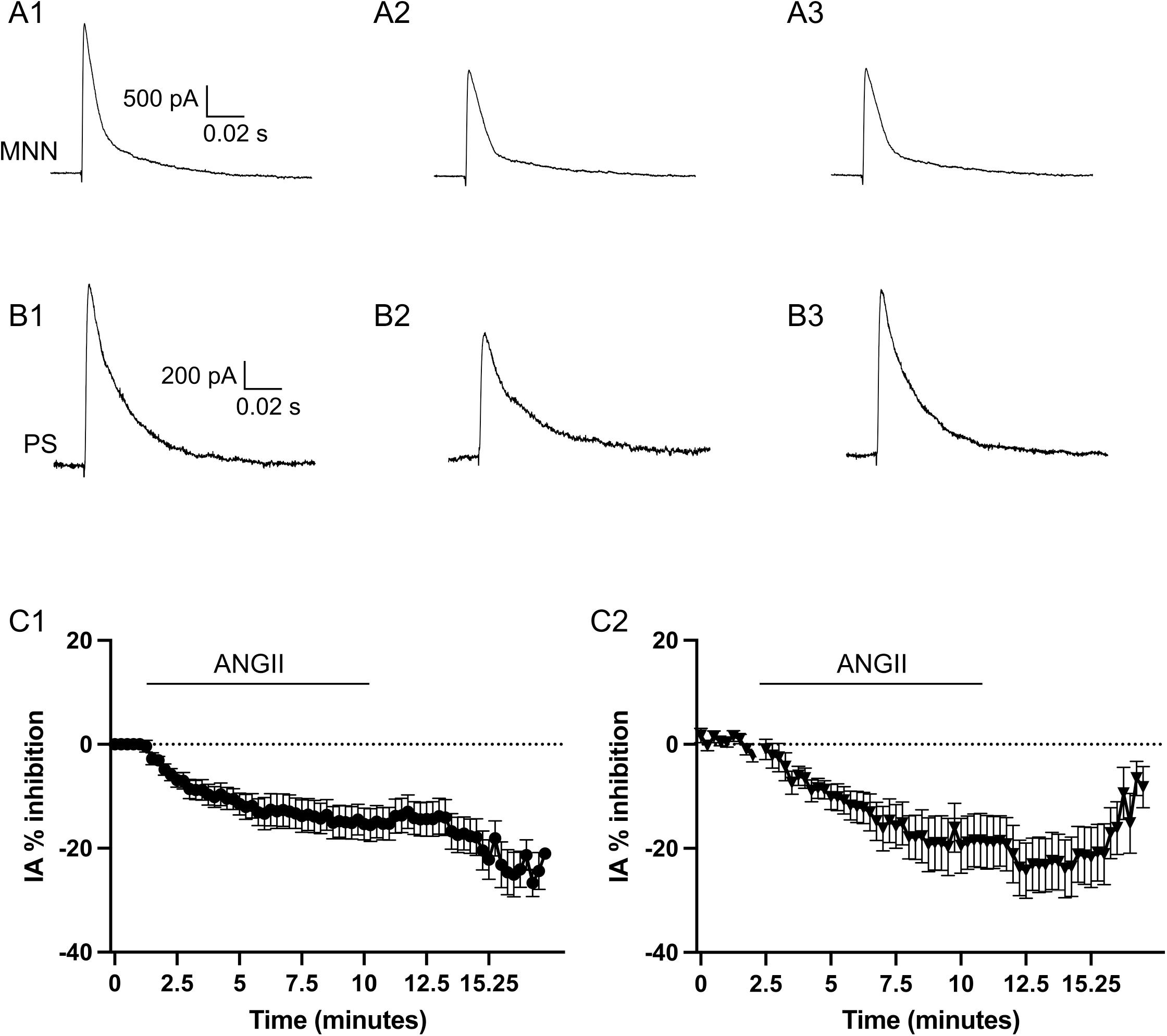
Angiotensin II (ANG II) inhibits I_A_ in MNNs and PSs neurons in the PVN. **A,** Representative traces of I_A_ in a MNN before (A1), during (A2), and after (A3) bath application of ANGII (0.5 μM, 8 mins.). **B**, Representative traces of I_A_ in a PS neuron before (B1), during (B2), and after (B3) bath application of ANGII (0.5 μM, 8 mins.). **C**, plots of percent I_A_ inhibition induced by ANGII over time in MNNs (C1, n= 15) and PSs (C2, n= 19).

The decay time course of the evoked I_A_ was best fitted by a two-exponential function. ANGII did not affect the decay kinetics of I_A_ in MNNs (τ1 control: 7.03 ± 1.38 ms; τ1 ANGII: 11.05 ± 2.30 p= 0.2; τ2 control: 57.45 ± 6.34 ms; τ2 ANGII: 74.4 ± 0.23 ms p= 0.1) or in PSs (τ1 control: 7.46 ± 1.96 ms; τ1 ANGII: 6.78 ± 1.76 p= 0.79; τ2 control: 61.51 ± 10.42 ms; τ2 ANGII: 57.70 ± 9.97 ms p= 0.79).

### ANGII-mediated inhibition of I_A_ is not dependent on glutamate signaling

We recently showed that ANGII in the PVN inhibits glutamate transporter activity, resulting in the buildup of extracellular glutamate (22) which, by acting on extrasynaptic NMDARs, could lead to inhibition of I_A_ (43). Thus, to determine whether ANGII-mediated I_A_ inhibition was mediated by glutamate, we repeated a set of experiments in the presence of the broad-spectrum glutamate ionotropic receptor blocker Kynurenic acid (KYN, 1 mM). As shown in ***Fig.2***, ANGII-mediated inhibition of I_A_ persisted in the presence of KYN both in MNNs (p= 0.53 vs ANGII-control, n=8) and in PSs (p= 0.39 vs ANGII-control, n=7).

**Fig 2.**
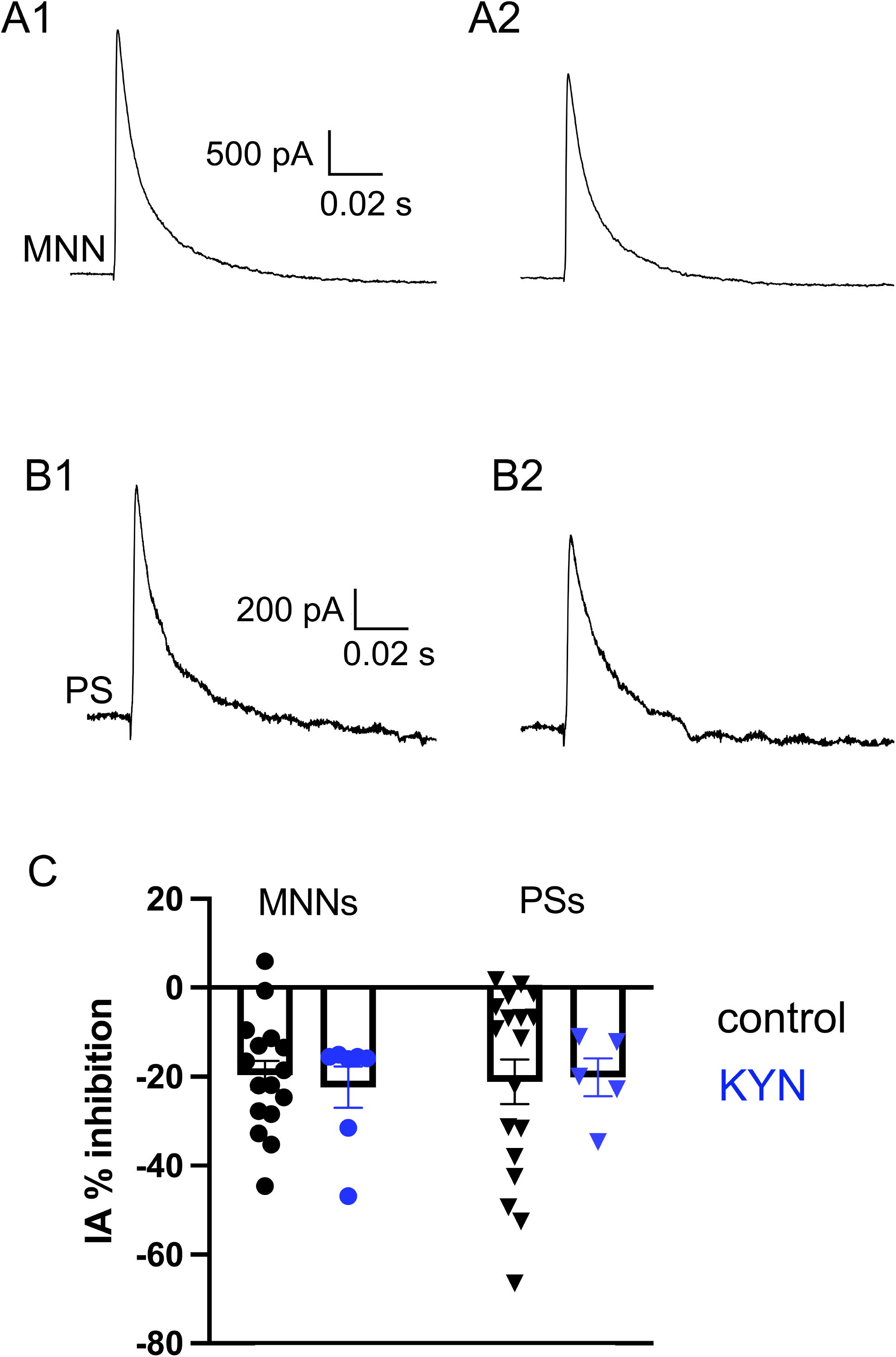
ANGII-mediated inhibition of I_A_ persists in the presence of ionotropic glutamate receptor blockade. **A,** Representative traces of I_A_ in a MNN before (A1) and during (A2) bath application of ANGII (0.5 μM, 8 mins.) in the presence of Kynurenic acid (KYN, 1 mM). **B**, Representative traces of I_A_ in a PS neuron before (B1) and during (B2) bath application of ANGII (0.5 μM, 8 mins.) in the presence of Kynurenic acid (KYN, 1 mM). **C**, Bar graph summarizing the mean ANGII-evoked inhibition of I_A_ in the presence of control ACSF (black) or in KYN (blue) MNNs (n= 15 and 8, respectively) and in PSs (n= 19 and 7, respectively).

### ANGII-mediated inhibition of I_A_ is also observed in PVN neurons of HF rats

The basal magnitude of I_A_ current density did not differ between sham and HF rats, both for MNNs (sham: 81.6 ± 7.9 pF; HF: 92.8 ± 9.5 pF, n= 29 and 21, respectively, p= 0.4, unpaired t test) and PSs (sham: 14.3 ± 0.9 pF; HF: 14.0 ± 1.0 pF, n= 28 and 9, respectively, p= 0.9, unpaired t test). Similarly, I_A_ decay kinetics were not altered in HF, when compared to sham rats (τ1: 6.73 ± 1.85 ms; τ2: 43.47 ± 6.82 ms; p= 0.91 and p= 0.22, respectively vs sham rats (see values above).

Similar to sham rats, we found ANGII to inhibit I_A_ both in MNNs (F= 7.02, p< 0.0001, One way ANOVA, n= 8) as well as in PSs (F= 2.85, p< 0.0001, One way ANOVA, n= 8) (***Fig.3***). No differences in the peak of the ANGII-mediated inhibition of I_A_ were observed between sham and HF rats in MNNs (p= 0.33, unpaired t test) or PSs (p= 0.63, unpaired t test).

**Fig 3.**
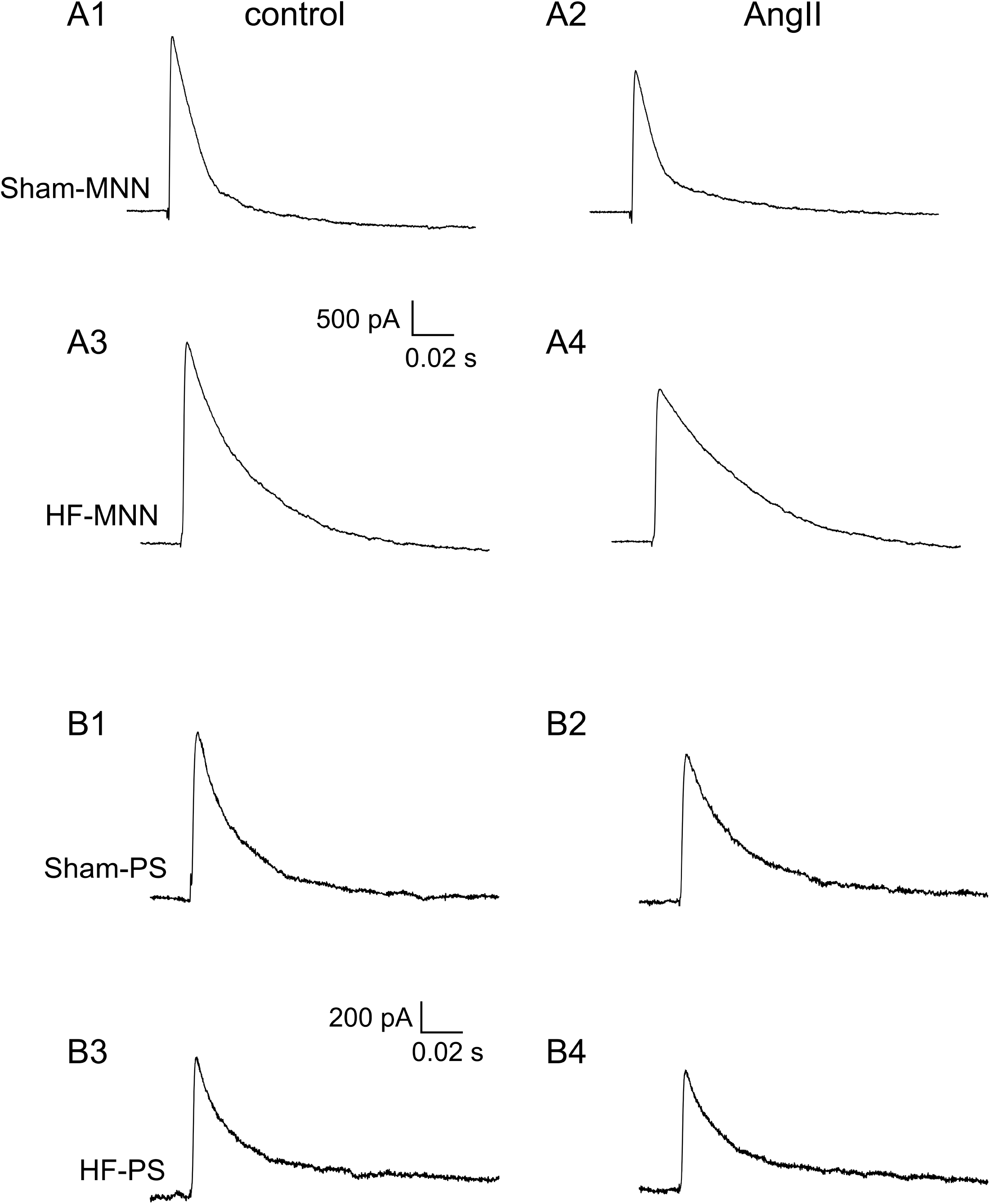
ANGII-mediated inhibition of I_A_ is observed both in sham and HF rats. **A,** Representative examples of ANGII-mediated inhibition of I_A_ in MNNs in a sham rat (A1,A2) and in a HF rat (A3,A4). A1 and A3: baseline traces; A2 and A4: in ANGII (0.5 μM). **B,** Representative examples of ANGII-mediated inhibition of I_A_ in PSs in a sham rat (B1,B2) and in a HF rat (B3,B4). B1 and B3: baseline traces; B2 and B4: in ANGII (0.5 μM).

### ANGII-mediated inhibition of I_A_ involves the MAPK-ERK intracellular pathway

To determine the contribution of the MAPK-ERK pathway to ANGII-mediated inhibition of I_A_, we performed a first set of experiments in which slices were preincubated in the presence or absence of the MAPK-ERK blocker PD98059 (50 μM, 1 h). Representative trace samples are shown in ***Fig.4*** while the mean summary data is shown in ***Fig.5***. We found that preincubating slices in PD98059 almost eliminated the ANGII-mediated inhibition of I_A_ in MNNs and PSs in sham rats (F= 0.69, p= 0.97, n= 9 and F= 082, p= 0.84, n= 13, respectively). Moreover, the mean peak ANGII-mediated I_A_ inhibition in the presence of PD98059 was significantly smaller than that observed in ANGII (p< 0.001 for MNNs and p< 0.002 for PSs, see ***Fig.5***).

**Fig 4.**
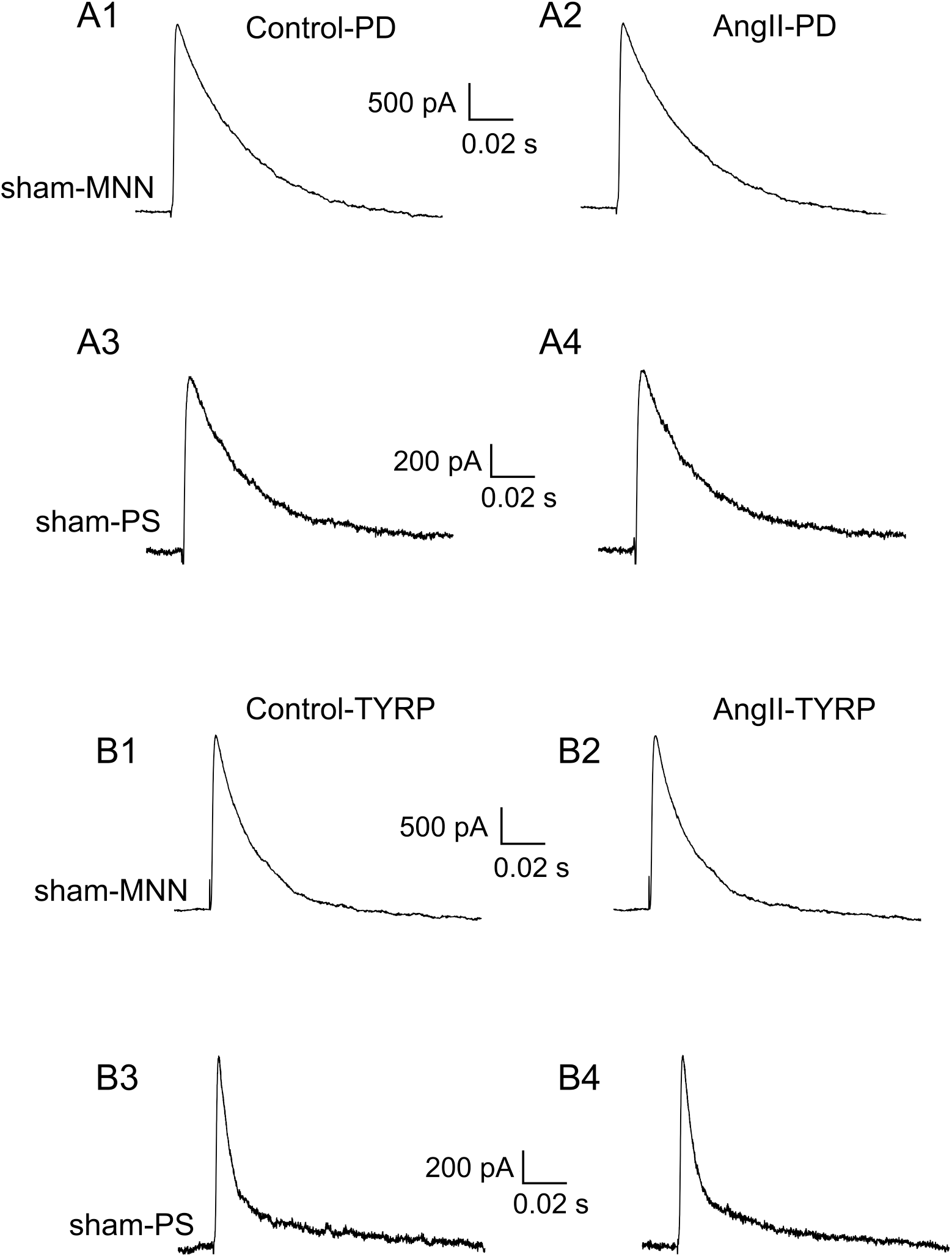
ANGII-mediated inhibition of I_A_ is prevented by MAPK-ERK and EGFR blockade. **A,** Representative examples showing lack of ANGII effects on I_A_ in the presence of the MAPK-ERK blocker PD98059 (PD, 50 μM) in a MNN (A1, A2) and in a PS neuron (A3, A4). **B,** Representative examples showing lack of ANGII effects on I_A_ in the presence of the EGFR blocker Tyrphostin (TYRP, 50μM) in a MNN (B1, B2) and in a PS neuron (B3, B4). All the traces were obtained from sham rats.

**Fig 5.**
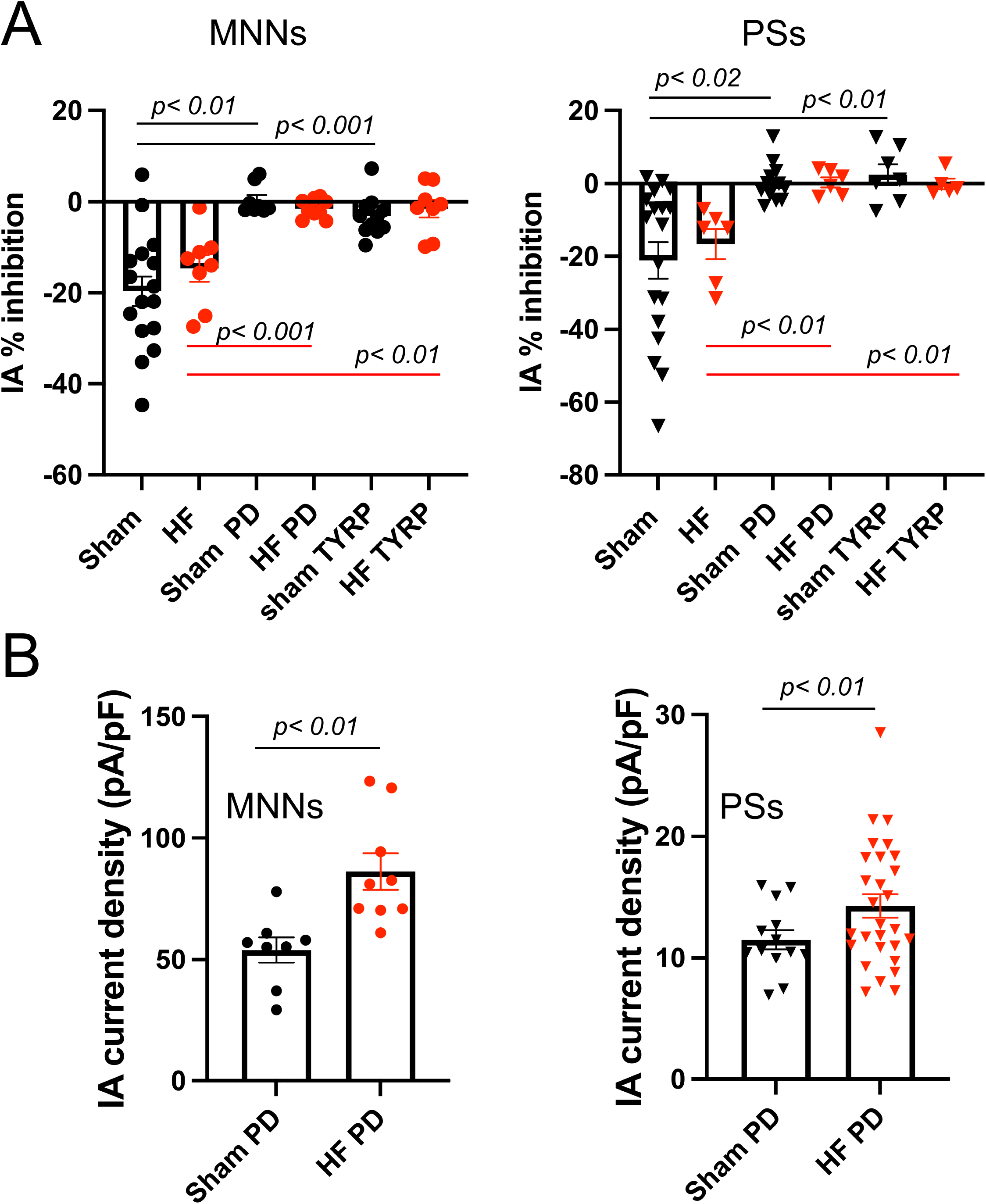
Summary of ANGII-mediated effects on I_A_ in control conditions and in the presence of MAPK-ERK and EGFR blockade. **A,** Bar graph summarizing the mean ANGII-evoked inhibition of I_A_ in MNNs (left) and PSs (right) in sham and HF rats in control conditions, and in the presence of the MAPK-ERK blocker PD98059 (PD, 50 μM) or the EGFR blocker Tyrphostin (TYRP, 50μM). **B**, Bar graphs summarizing the mean I_A_ current density in the presence of PD98059 in sham and HF rats in MNNs (left) and PSs (right).

Similar effects were observed in HF rats. In MNNs, whereas ANGII effects on I_A_ were still found to be significant (F= 3.4 p< 0.001. n= 10), the mean peak ANGII-mediated I_A_ inhibition in the presence of PD98059 in MNNs was significantly smaller than that observed in ANGII control (p< 0.001). In PSs, the ANGII inhibition of I_A_ was almost completely blunted (F= 0.38, p= 0.99, n= 6). Similarly, the mean peak ANGII-mediated I_A_ inhibition in the presence of PD98059 in PSs was significantly smaller than that observed in ANGII control (p< 0.01, unpaired t test). These results strongly support the involvement of the MAPK-ERK pathway in mediating the inhibitory effect of ANGII on I_A_ in both PVN neuronal types, and in both sham and HF rats.

### ANGII-mediated inhibition of I_A_ involves EGFR signaling

Finally, in a separate set of experiments we sought to determine the contribution of the epidermal growth factor receptor (EGFR) to the ANGII-mediated effects on I_A_. EGFR is a receptor tyrosine kinase that may be transactivated by G-protein-coupled receptors (44) including ANGII (45), to induce MAPK-ERK signaling. Slices were preincubated in the presence or absence of the EGFR blocker Tyrphostin (50μM, 1 h). Representative trace samples are shown in ***Fig.4*** while the mean summary data is shown in ***Fig.5***. We found that preincubating slices in Tyrphostin in sham rats (F= 0.22, p> 0.99, n= 10 and F= 0.24, p> 0.99, n= 7, respectively) almost eliminated the ANGII-mediated inhibition of I_A_ MNNs and PSs respectively. Moreover, the mean peak ANGII-mediated I_A_ inhibition in the presence of Tyrphostin was significantly smaller than that observed in ANGII (p< 0.001 for MNNs and p< 0.01 for PSs, see ***Fig.5***).

Similarly, preincubating slices in Tyrphostin almost eliminated the ANGII-mediated inhibition of I_A_ in MNNs and PSs in HF rats (F= 1.33, p= 0.052, n= 9 and F= 0.55, p> 0.99, n= 5, respectively). Moreover, the mean peak ANGII-mediated I_A_ inhibition in the presence of Tyrphostin was significantly smaller than that observed in ANGII (p< 0.01 for both MNNs and PPSs, see ***Fig.5***). These results support the involvement of the EGFR pathway in mediating the inhibitory effect of ANGII on I_A_ in both PVN neuronal types and both in sham and HF rats.

Neither PD98059 nor Tyrphostin treatments significantly affected the baseline magnitude of I_A_ in MNNs (PD98059 sham: p= 0.20 unpaired t test; HF: p= 0.94 unpaired t test; Tyrphostin: sham: p= 0.54 unpaired t test; HF: p= 0.11 unpaired t test) or PSs neurons (PD98059 sham: p= 0.08 unpaired t test; HF: p= 0.25 unpaired t test; Tyrphostin: sham: p= 0.12 unpaired t test; HF: p= 0.90 unpaired t test). Despite this however, we did find that in slices preincubated with PD98059 the basal magnitude of I_A_ was significantly larger in HF compared to sham both in MNNs and PSs (p= 0.006 in both groups, unpaired t test) (***Fig.5***). This however, was not the case for slices preincubated with Tyrphostin (not shown). These results suggest a certain level of overactivation of the ANGII-MAPK-ERK pathway in PVN neurons of HF rats, leading to tonic inhibition of I_A_ in this condition.

## DISCUSSION

Several studies have indicated that a signaling mechanism by which ANGII acts at the cellular level involves the ubiquitous intracellular MAPK cascades (24, 25). Moreover, we showed in recent studies that ERK1/2 signaling is involved in ANGII-mediated sympathetic excitation within the PVN, and that inhibition of brain MAPK signaling significantly reduces sympathetic nerve activity in HF rats (13, 28). Still, the precise cellular mechanisms by which the ANGII-MAPK-ERK1/2 signaling cascade influences sympathohumoral outflow from the PVN remain to be determined. Given that the latter is directly dependent on the firing activity of PVN neurosecretory and presympathetic neurons, we obtained in the present study whole-cell patch-clamp recordings from identified PVN neurons, and assessed the effects of ANGII on the A-type K^+^ current (I_A_), a major intrinsic conductance that plays a key role in modulating the firing activity of both neuronal populations (34–36). Our main results could be summarized as follows: (1) ANGII inhibited I_A_ both in magnocellular neurosecretory and presympathetic PVN neurons; (2) the ANGII-mediated inhibition of I_A_ was almost completely blocked by a MAPK-ERK pathway blocker as well as an EGFR blocker; and (3) similar effects were observed in PVN neurons from sham and HF rats.

The A-type potassium current (I_A_) is a dominant intrinsic, voltage-dependent subthreshold current that determines interspike interval and firing discharge in both MNNs and presympathetic neurons (34–36). Previous studies have shown than pharmacological inhibition of I_A_ results in increased firing discharge and frequency in both neurosecretory and presympathetic SON and PVN neurons (34–36, 43). Moreover, downregulation of I_A_ availability contributes to increased hypothalamic neuronal excitability during hypertension (37, 46), and a similar mechanism has been previously reported in NTS neurons of spontaneously hypertensive rats (47). Thus, while not directly tested in the present study, it is reasonable to assume that the ANGII-mediated inhibition of I_A_ constitutes a key underlying mechanism by which ANGII leads to increased firing activity of both magnocellular neurosecretory and presympathetic neurons, and thus neurohumoral output from the PVN.

Importantly, in addition to regulating somatic firing discharge, I_A_ in hypothalamic neurons modulates dendritic excitability, and as we previously showed, I_A_ inhibition facilitates dendritic Ca^2+^ signaling and propagation (46). A rise in dendritic Ca^2+^ in MNNs results in the local release of oxytocin and vasopressin (48). Thus, by inhibiting I_A_ in VP neurons, ANGII likely results in enhanced dendritic Ca^2+^ signaling, potentiating in turn dendritic release of VP. This is functionally important because we recently showed than dendritically released VP in the PVN can diffuse in the extracellular space to recruit and stimulate the activity of presympathetic neurons, further increasing neurohumoral outflow from the PVN (49).

Whether ANGII effects on neurosecretory and presympathetic PVN neurons (including the inhibition of I_A_ reported here) is the result of direct actions of ANGII on these neurons, or alternatively, whether it involves other intermediaries, remains a somewhat controversial issue. In this sense, recent studies indicate that ANGII AT1 receptors in the PVN are not expressed in these neurons, but rather their expression is limited to CRH and TRH neurosecretory neurons (23, 50). In fact, we recently showed that activation of AT1R-expressing CRH neurons in the PVN leads to a CRFR1-receptor mediated stimulation of presympathetic neurons and sympathetic outflow from the PVN (23), supporting a functional crosstalk between CRH and presympathetic neurons in the PVN. Moreover, we previously showed that AT1 receptors are also expressed in PVN astrocytes, and that their activation inhibits astrocyte glutamate transporter activity, leading to a buildup of extracellular glutamate and activation of extrasynaptic NMDA receptors in presympathetic PVN neurons (22). Our current experiments showing that ANGII-mediated inhibition of I_A_ persisted in the presence of glutamate receptor blockade argues against this indirect pathway in mediating these effects. Still, we cannot conclusively determine yet whether other intermediaries (e.g., CRF) are involved.

Notable in this regard is a recent report that sympathetic excitation induced by PVN microinjection of TNF-a does appear to be mediated by glutamate (51). TNF-a, like ANGII, induces ERK1/2 signaling and related genomic neurochemical changes in the PVN(29), but the effect of TNF-a-induced ERK1/2 signaling on I_A_ in PVN neurons has not been tested. However, the failure of glutamate receptor blockade to influence ANGII-induced decreases in I_A_ suggests differences in the downstream signaling pathways mediating the excitatory responses to RAS and neuroinflammation in HF.

One pathway than leads to ERK1/2 signaling activation is stimulation of EGFR (52). EGFR is not only activated by its endogenous ligands, i.e., the epidermal growth factor and the transforming growth factor-alpha (53), but is also activated or transactivated by ANGII and proinflammatory cytokines (54–58). In fact, we recently showed that EGFR contributes to increased ERK1/2 signaling and augmented sympathetic excitation in HF (59). That study suggested an effect of EGFR-induced ERK1/2 signaling on neurochemical mediators of sympathetic excitation in the PVN in HF. The present study, demonstrating similar effects of an EGFR blocker and an ERK1/2 pathway blocker to prevent the ANGII-induced reduction in I_A_ in the PVN of sham and HF rats, suggests a prominent role for the EGFR in the ANGII-induced increases in PVN neuronal excitability.

Rats with ischemic HF are characterized by having an exacerbated sympathohumoral activation (4–7, 12) and increased PVN neuronal excitability (4, 38, 60–63). Importantly, and as stated above, overactivation of the RAS, acting in part via the MAPK-ERK1/2 cascade (27, 28), has been proposed as a key signaling mechanism contributing to sympathetic output in this disease. Given this, and our results showing that ANGII inhibits I_A_ via the MAPK-ERK1/2 pathway, we speculated to observe a basally blunted I_A_ availability in PVN neurons in HF rats as we previously reported in hypertensive conditions (37, 46). While we did not find a diminished I_A_ magnitude in HF when compared to sham rats under control conditions as expected, we did find than that I_A_ magnitude was significantly larger in both PVN neuronal types in HF rats after blockade of MAPK with PD98059. These results support the notion that in HF, I_A_ is tonically inhibited to a larger extent via activation of the MAPK cascade. Still, whether this is due to an increased ANGII signaling during HF, or due to other signals that converge onto this similar cascade, including aldosterone, perhaps via an effect on ANGII AT1 receptors (64) and proinflammatory cytokines (29), remains at present unknown. Future studies aiming at determining (a) the precise mechanisms underlying differential modulation of I_A_ in HF, and (b) the contribution of I_A_ to exacerbated neurohumoral outflow in this condition are warranted.

## PERSPECTIVES

Recent studies have demonstrated that MAPK ERK1/2 signaling contributes to the exaggerated sympathohumoral activity in HF by upregulating RAS activity and inflammation in the PVN, chronic genomic influences of MAPK-ERK requiring gene transcription. In that capacity, MAPK-ERK signaling stimulates the production of many of the same agents that elicit its activity, perpetuating the HF condition. The present study identifies a more immediate direct cellular effect of MAPK-ERK signaling, influencing the activity of a key K^+^ current (I_A_) that normally regulates the excitability of presympathetic neurons that drive sympathetic nerve activity and of neuroendocrine neurons that regulate VP release. While the present study focused on the effects of ANGII on IA and membrane excitability of these PVN neuronal populations, these findings may well apply to other excitatory agents – e.g., pro-inflammatory cytokines – that activate MAPK-ERK signaling, and to other settings – e.g., hypertension – in which brain RAS activity and neuroinflammation mechanisms appear to be contributing to disease progression.

